# Maternal Anxious Attachment Style is Associated with Reduced Mother-Child Brain-to-Brain Synchrony During Passive TV Viewing

**DOI:** 10.1101/2020.01.23.917641

**Authors:** Atiqah Azhari, Giulio Gabrieli, Andrea Bizzego, Marc H. Bornstein, Gianluca Esposito

## Abstract

Synchrony in developmental science reflects the coordination of mother and child to the same mental state. Mentalisation processes are influenced by individual attachment styles. A mother with an anxious-related attachment style tends to engage in emotional mentalisation that relies on her child’s social cues. During an everyday joint activity of watching television shows together, we hypothesised that anxiously-attached mothers are less able to match their mental state to characters in the shows as their attention is likely detracted from the show and directed towards the child. We predict that this mismatch in mother’s and child’s emotional states would be reflected in reduced dyadic brain-to-brain synchrony. To test this hypothesis, we profiled mothers’ Maternal Anxiety score using the Preoccupation and Need for Approval subscales of the Attachment Style Questionnaire (ASQ) and used functional Near-infrared Spectroscopy (fNIRS) hyperscanning with 33 mother-child dyads to measure prefrontal cortex (PFC) synchrony while the dyads watched three 1-min animation videos together. Greater Maternal Anxiety is associated with less synchrony in the medial right prefrontal cluster implicated in mentalisation processes. Anxiously-attached mothers appear to exhibit less brain-to-brain synchrony with their child which suggests differences in intersubjective shared experiences that potentially undermines the quality of bonding during everyday joint activities.

## Introduction

Adult attachment is the stability with which individuals cultivate emotional proximity with attachment figures in adulthood, the principal of which is one’s romantic partner (Hazan & Shaver, 1987). Profiles of adult attachment are broadly classified into secure and insecure (anxious or avoidant) styles (see Mikulincer & Shaver (2010) for an overview). The Attachment Style Questionnaire (ASQ) is a commonly used instrument to categorise adult attachment styles with theoretical underpinning in a five-factor solution: Confidence (in self and others), Discomfort with Closeness, Relationships as Secondary, Need for Approval and Preoccupation with Relationships (Feeney & Noller, 1991). While Confidence certainly connotes secure attachment, each of the other four factors reflect a distinctive facet of insecure attachment. Discomfort with Closeness serves as a central factor for Hazan & Shaver (1987) model of avoidant attachment. Analogously, Relationships as Secondary is contingent with Bartholomew (1990) conceptualisation of a dismissing attachment style, in which individuals assert independence and autonomy as a barrier to shield their vulnerability from emotional pain. Both Discomfort with Closeness and Relationships as Secondary are factors that form the avoidant attachment style (Sperling & Berman, 1994). The last two factors, Need for Approval and Preoccupation with Relationships, characterises the anxious attachment style, in which the former reflects individuals’ needs for confirmation and acceptance from others (Bartholomew & Horowitz, 1991), and the latter describes individuals who approach others so as to meet their needs for dependency (Hazan & Shaver, 1987).

Maternal adult attachment styles predict the quality of their caregiving behaviours (Selcuk et al., 2010). Mothers who are insecurely attached may experience difficulties in regulating her emotional state to respond optimally to the needs of her child (Jones et al., 2014). Those with insecure attachment have a greater tendency of being insensitive (Jones et al., 2018; van IJzendoorn, 1995) and emotionally unavailable (Licata et al., 2016a), and are more likely to engage in harsh parenting practices (Jones et al., 2014). In a study by Selcuk et al. (2010), the authors observed mother-child interactions and investigated whether caregiving behaviours were associated with caregiving practices. Congruent with several other studies (Edelstein et al., 2004; Rholes et al., 1995), they found avoidant attachment to be associated with diminished maternal sensitivity during mother-child interactions. Since anxious mothers desire extreme proximity with their children (Buehler, 2017), it was unexpected that anxious attachment was correlated with missing the child’s signals. The authors reasoned that mothers with anxious attachment maintain closeness to allay their persistent worries about being accepted, thereby fulfilling their own attachment needs, rather than that of their child (Mikulincer & Shaver, 2007). This myopic focus may hinder the mother’s ability to detect and accurately interpret her child’s signals during dyadic interactions (Feeney & Collins, 2001). Regrettably, the maladaptive parenting practices that accompany maternal insecure attachment styles reduce the quality of interaction in a mother-child dyadic relationship (Licata et al., 2016b).

Synchrony refers to the reciprocal and congruent exchange of behavioural and physiological signals between dyadic partners (Fleming et al., 1999; Ruth Feldman, 2012). Over repeated experiences with each other, a mother and child become accustomed to each other’s emotional signals and behavioural repertoire. In a mother-child relationship, such attunement to each other’s social cues leads to synchrony, which manifests at the behavioural level in the form of matched gazes, facial expressions and vocalisations (Feldman, 2012; Feldman, 2007). Since accurately perceiving and responding to a child’s signals is a bedrock for mother-child synchrony, mothers with anxious attachment, who have been found to overlook their child’s signals (Selcuk et al., 2010), might have difficulties establishing synchrony. Selcuk et al. (2010) also found that mothers with avoidant attachment exhibited non-synchronous interactions with their children due to discomfort with close contact, in addition to missing the child’s signals. In an earlier study, Crandell et al., (1997) showed that secure mothers engaged in interaction to a higher degree of synchrony compared to their insecure counterparts. These findings cumulatively suggest that maternal attachment influences mother-child synchrony at the behavioural level.

Beyond behavioural signals, synchrony has been shown to exist at the physiological level. In fact, synchrony at the behavioural level is postulated to be facilitated by entrainment of biological signals, including the matching of brain signals between dyadic partners (e.g. Behrendt et al., 2020; Nguyen et al., 2020; Piazza et al., 2020; Reindl et al., 2018). Employing hyperscanning techniques, where multiple brains are simultaneously recorded in tandem, Piazza et al. (2020) demonstrated that adult and infant brains synchronise with each other when engaging in a natural interaction. Brain-to-brain synchrony specific to the parent-child dyad has also emerged. For example, Reindl et al. (2018) designed a study where mother-child and stranger-child dyads engaged in a cooperative and competitive game with each other. Synchrony emerged in several regions in the prefrontal cortex (PFC), including the dorsolateral PFC and the frontopolar cortex, only in the mother-child dyad during the cooperative game. In another study, Nguyen et al. (2020) tasked mothers and children to solve a tangram puzzle either individually or together. The researchers found greater synchrony in the bilateral PFC and the temporal-parietal regions when dyads solved the puzzle together. Moreover, neural synchrony was also accompanied by higher incidences of behavioural synchrony, further cementing the notion of physiological entrainment that is unique to the parent-child dyad.

Across many studies, one of the most commonly implicated regions in brain-to-brain synchrony is the PFC (Azhari et al., 2019; Azhari et al., 2020; Behrendt et al., 2020; Nguyen et al., 2020; Reindl et al., 2018). The PFC is the seat of the mutual social attention system, which facilitates shared attention between people (Gvirts & Perlmutter, 2020). Mutual attention is well-established to be the foundation of social cognition (Tomasello, 1995). The PFC has been associated with both attention processing (e.g., Hopf & Mangun, 2000)) and theory of mind (Shamay-Tsoory & Aharon-Peretz, 2007), both of which fall into the same network that oversees mentalisation processes (Andrews-Hanna et al., 2014). Mentalisation processes in the PFC are also linked to attachment style. According to a model by (Fonagy & Luyten, 2009; Lieberman, 2007)), mentalisation can be distinguished into two processes that are adopted differentially based on adult attachment style: one is emotional and the other is cognitive mentalisation. Emotional mentalisation emphasises automatic and implicit processing of others’ emotions based on some concrete physical attributes, such as facial expressions. In contrast, cognitive mentalisation involves voluntary and conscious processing of other’s emotions by inferring some internal attributes, such as feelings and intentions (Fonagy & Luyten, 2009; Lieberman, 2007). Mentalisation is mediated by an extensive brain network, including the lateral PFC (Fonagy & Luyten, 2009). Individual attachment styles may influence cognitive and emotional mentalisation strategies. For instance, an anxious attachment style promotes hypervigilance to signals of social rejection, thereby enhancing emotional mentalisation (Fonagy & Luyten, 2009). From these studies, we have ascertained that mentalisation processes that reside in the PFC serve as a basis for both mutual attention during social engagement, and have been shown to be influenced by attachment styles.

Till date, no study has investigated the role of maternal attachment style on mother-child brain-to-brain synchrony during passive engagement. The present study is an extension of a study by (Azhari et al., 2019) who examined mother-child brain-to-brain synchrony in relation to parenting stress using tandem functional near-infrared spectroscopy (fNIRS). fNIRS records changes in local blood oxygenation such that higher levels of oxygenated blood serve as a proxy of brain activation. Greater parenting stress was associated with diminished synchrony in the dyad’s medial left cluster of the PFC when the pair engaged in an everyday activity of watching animation videos together. Findings from (Azhari et al., 2019) suggest the moderating effect of maternal variables on brain-to-brain synchrony during passive joint viewing of animation shows, which probes further investigation as to whether maternal attachment style exerts an effect of brain-to-brain synchrony too. While the vast majority of studies have investigated synchrony during active social interaction, there are two main reasons to expect brain-to-brain synchrony to occur during passive joint animation watching too. First, comprehension of a movie’s plot necessitates social cognitive processes such as understanding the perspective of the characters in the show and interpreting their verbal and non-verbal signals (Lyons et al., 2020), all of which are facilitated by mentalisation regions in the PFC (Yeshurun et al., 2017). Second, in a seminal paper by (Hasson et al., 2004), the authors found that people’s brains tend to show similar activations during natural viewing of movie scenes, even in the absence of direct communication. Inter-subject synchrony was found well beyond visuo-spatial brain regions, and was observed in higher-order cortical regions. Applying these findings to the present study, it is thus probable that brain-to-brain synchrony would emerge in areas responsible for mentalisation processes in mother-child dyads.

We embarked on this study with a central hypothesis. We predicted that less brain-to-brain synchrony in the medial PFC regions, responsible for mentalisation, would be observed in dyads in which the mother has a higher anxiety-related attachment style. In line with the mentalisation models proposed by (Fonagy & Luyten, 2009; Lieberman, 2007), anxiously-attached mothers whose strategy is to employ emotional mentalisation and rely on concrete social cues, are more likely to be preoccupied with being accepted by their child. This fixation may interfere with the mothers’ ability to attend to the animation stimuli and process the plot and characters screened to them, thus impeding brain-to-brain synchrony. Overall, findings from this study would lend significant insight into how maternal attachment style influences joint processing of narrative scenes which mother-child dyads commonly engage in when watching television shows together at home.

## Methods

### Participants

In total, 33 mother-child dyads (14 girls) (Mean age of mothers=34.7 years, S.D.= 4.1 years; Mean age of children=41.9 months, S.D.= 6.1 months) participated. Participants were recruited through online platforms (forums and Facebook groups) before they were screened for cognitive, visual, hearing, or developmental impairments that might hinder them from comprehending the task stimuli. Participants signed Informed consents, and each dyad received monetary compensation upon completion of the study. This study was approved by the Institutional Review Board of XXX University.

### Experimental Procedure

Mothers first completed an attachment style questionnaire. Then, mothers’ and children’s brain responses were investigated using tandem functional near-infrared spectroscopy (fNIRS) while they watched three animation videos. All videos were presented on a laptop approximately 40 cm in front of the dyad who were alone in the test room (Figure 1 from Azhari et al. 2019).

**Figure 1.**
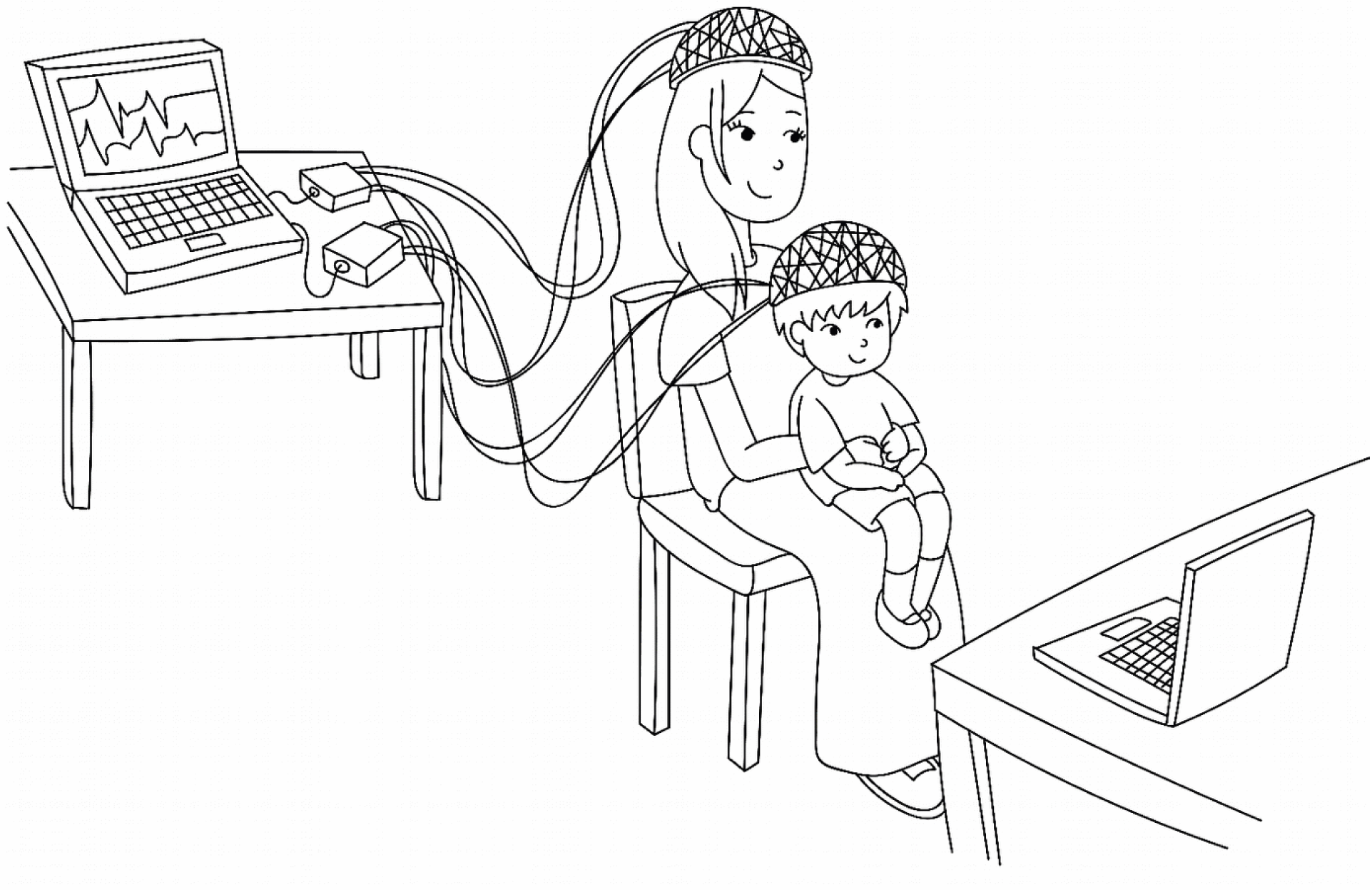
Experimental arrangement during the fNIRS session.

### Questionnaire

The Attachment Style Questionnaire (ASQ; Feeney et al., 2014) is a 40-item instrument that measures an adult’s attachment style in regard to general (rather than specific romantic) relationships. The questionnaire employs a 6-point Lickert scale on which participants rate themselves and others. Items are grouped into five subscales: *Confidence, Discomfort with closeness, Relationships as secondary, Need for approval*, and *Preoccupation with relationships*. These subscales are grounded in classification theories of attachment styles, where *Need for approval* and *Preoccupation with relationships* indicate anxious attachment and *Discomfort with closeness* and *Relationships as secondary* reflect avoidant attachment (Stein et al., 2002). *Confidence* is related to secure attachment. Our main interest and hypothesis concerned scales having to do with anxious attachment, so we used the ASQ to derive anxious attachment status in mothers.

### Video Stimulus

Participants were presented with three 1-min video clips that were excerpted from The Incredibles, Peppa Pig, and Brave. To increase the generalisability of the dyadic activity across different animation shows, the clips differed in emotional valences and visual and audio complexities. A 5-sec fixation cross appeared before the onset of the first clip, after which a 10-sec fixation cross appeared between each clip (Figure 2 from Azhari et al. (2019). The clips were presented in a semi-randomised manner on a 15-inch Acer Laptop. Both brightness and volume were set to 60%.

**Figure 2.**
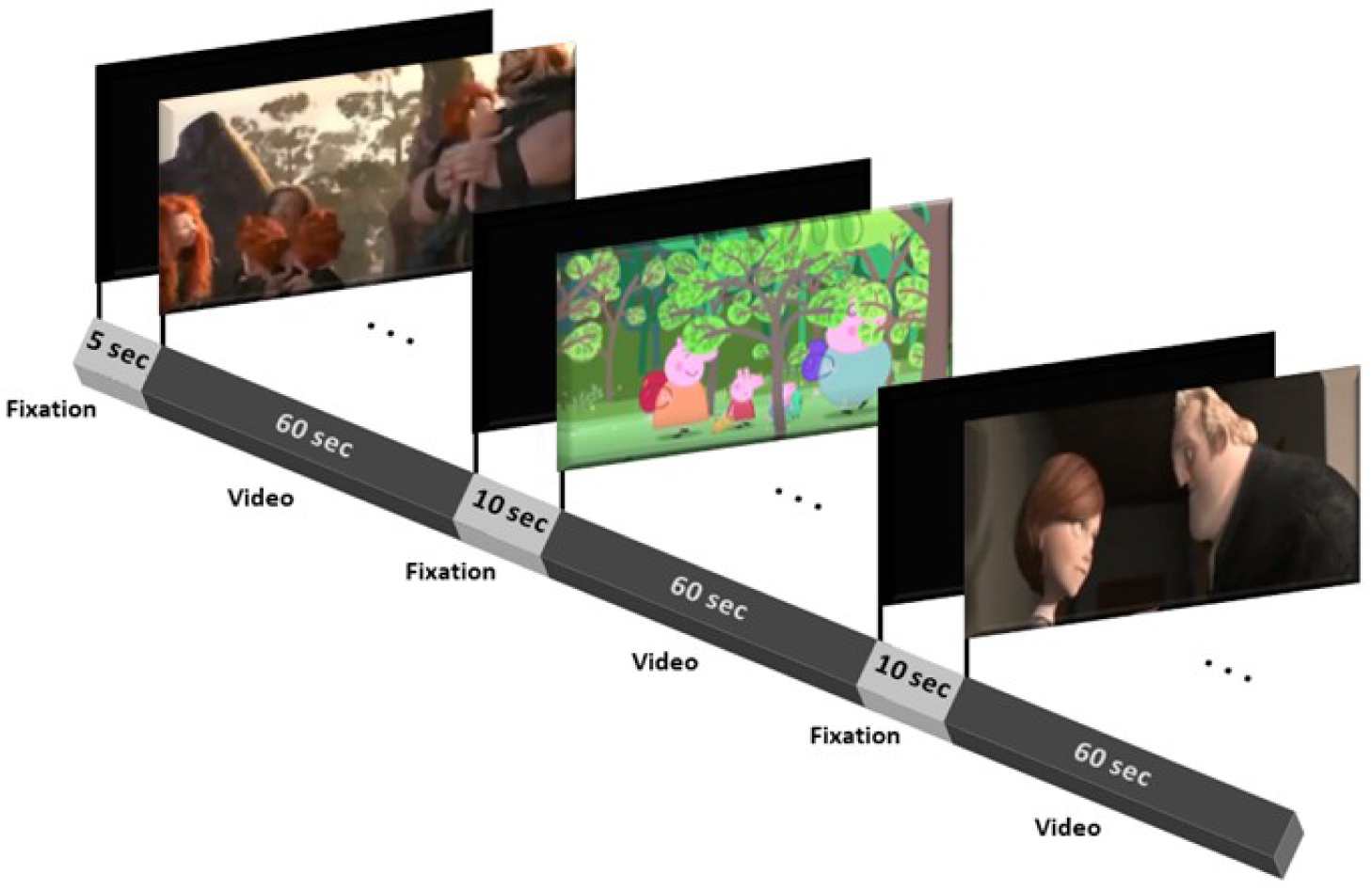
Diagram depicting the stimuli presentation. A fixation cross was screened for 5 sec before the onset of the first clip. A total of three 1-min video clips were presented with an inter-stimulus interval of 10 sec between each clip.

The visual complexity of each clip was quantified using Python and FFmpeg (v. 3.4.4). To analyse audio complexity, each video was first converted to an audio file on FFmpeg. Praat software (v. 6.0.46) was then used to extract audio intensity and fundamentals from the audio files. To compute the emotional valence of each clip, two independent coders rated each sec of the clip as either positive, neutral, or negative. The sum of the sec-by-sec coding across the entire clip was used to rate positivity. Table 1 shows the results of visual and audio complexities and emotional valences of each clip.

**Table 1.**
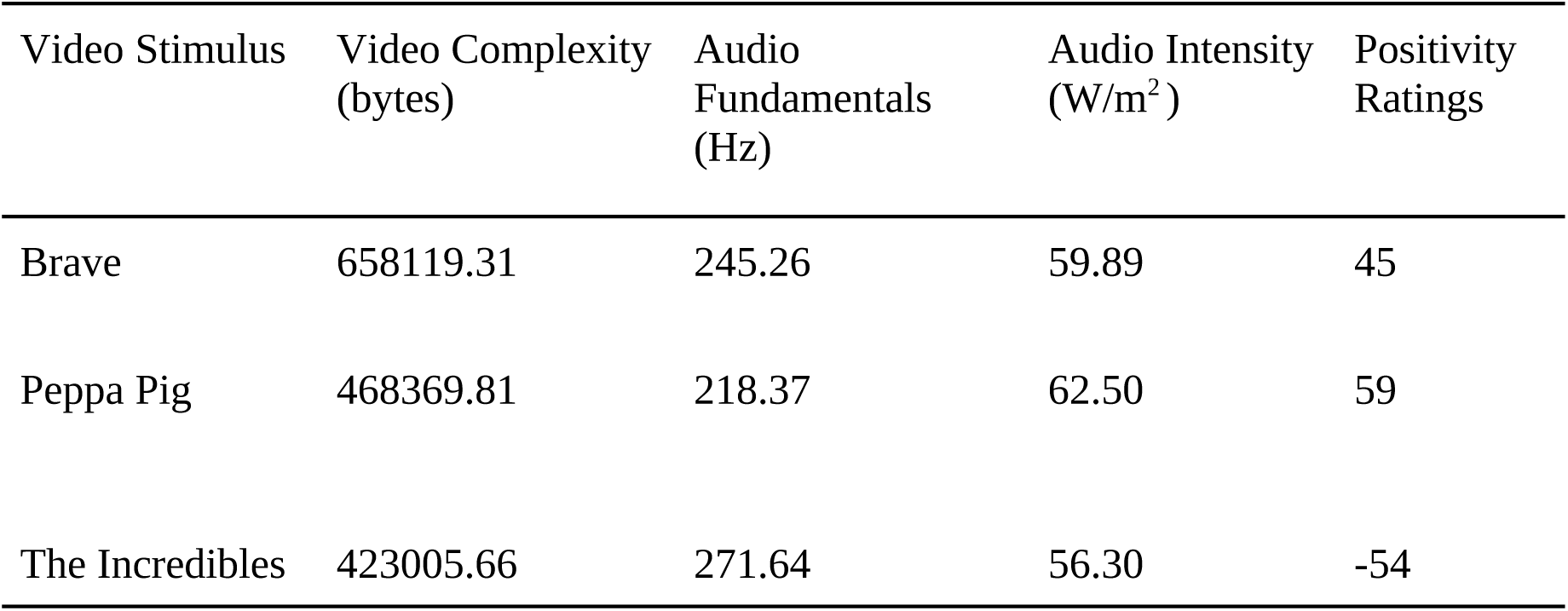
Video complexity, audio fundamentals, audio intensity, and positivity ratings of the three video stimuli excerpted from Brave, Peppa Pig, and The Incredibles.

### Functional Near-Infrared Spectroscopy (fNIRS)

#### fNIRS Data Acquisition and Pre-Processing

fNIRS recording was conducted in tandem hyperscanning mode at a scan rate of 7.81Hz (NIRSport, NIRx Medical Technologies LLC). 8 LED sources (wavelengths of 760nm and 850nm) and 7 detectors were arranged on the caps according to a standard PFC montage to form 20 channels on each cap (NIRS v.205 software). Distances between source and detector pairs were under an optimal maximum of 3 cm.

fNIRS pre-processing was done on the NIRSLab software (NIRS v.205 software). Markers that denote the onset of each video stimulus were added to the time series signals. Upon visual inspection of the signal, discontinuities and spike artefacts were manually removed. The 20 channels were then inspected for their gain and coefficient of variation (CV) scores which indicated the noise level in the signal. Channels with a gain>8 and CV>7.5 were deemed to have significant noise and subsequently discarded. A band-pass filter of 0.01-0.2 Hz was applied on the remaining channels to eliminate baseline shift variations and slow signals. To compute hemoglobin levels for each channel, pre-processed signals were converted to changes in concentration of oxygenated (HbO) and deoxygenated hemoglobin (HbR) as a function of the modified Beer-Lambert law. The resulting haemodynamic signals were visually examined by two independent coders to inspect for any further artifacts. Any artifacts detected by the coders were removed.

As there were 33 mother-child dyads, the total number of channels at the beginning was 66 subjects * 3 videos * 20 channels = 3960. Out of this total, 1283 channels were discarded (32.4%).

#### Cluster Grouping of Channels

As proximal regions of the PFC function in an interdependent manner, examining prefrontal responses in clusters rather than single channels lends a practical interpretation of the findings. Similar to Azhari et al. (2019), the channels were divided into four groups based on their physical proximity to each other to form the frontal left, frontal right, medial left, and medial right clusters as reported in Table 2. The corresponding Brodmann areas for each cluster are depicted in Figure 3.

**Table 2.** Control and copresence synchrony values in the four PFC clusters, along with the corresponding sample size.

|  |  | **Control** |  | **Copresence** |  |  |  |
| --- | --- | --- | --- | --- | --- | --- | --- |
| **Cluster** | **N** | **Mean** | **SD** | **Mean** | **SD** | **U** | **p** |
| Medial Left | 71 | 0.158 | 0.260 | 0.169 | 0.306 | 29458.0 | 0.333 |
| Medial Right | 68 | 0.145 | 0.268 | 0.189 | 0.253 | 27658.0 | 0.078 |
| Frontal Left | 95 | 0.156 | 0.255 | 0.200 | 0.264 | 77947.0 | 0.019 |
| Frontal Right | 92 | 0.151 | 0.266 | 0.169 | 0.277 | 64792.0 | 0.304 |
| **All** | **326** | **0.153** | **0.262** | **0.182** | **0.275** | **763629.0** | **0.013** |

**Figure 3.**
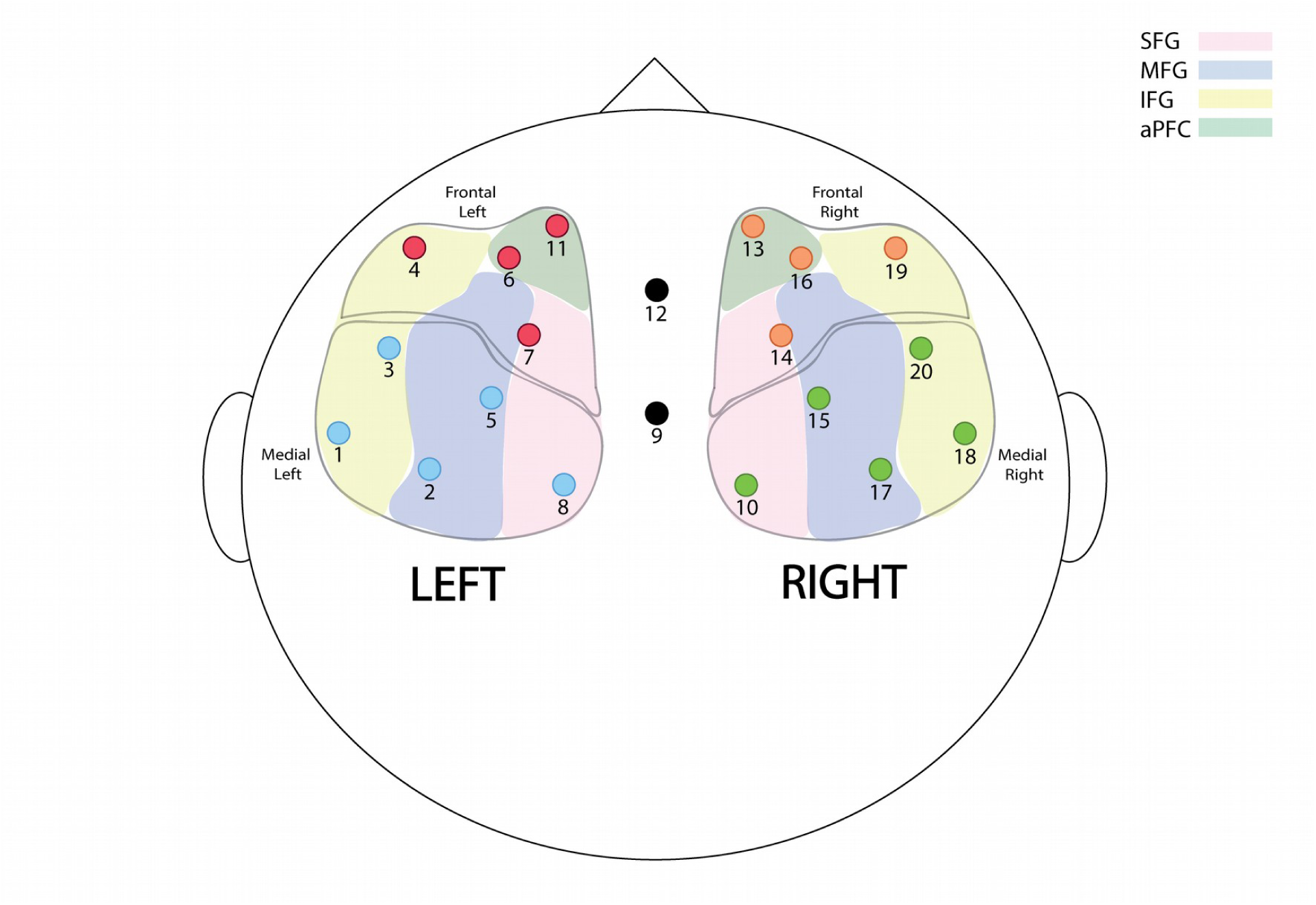
Positions of the 20 channels according to a standard prefrontal cortex montage. Note: SFG = superior frontal gyrus, MFG = middle frontal gyrus, IFG = inferior frontal gyrus, aPFC = anterior prefrontal cortex.

For each subject, we derived the signals associated with the brain activity of each cluster, by computing the average of the normalized signals of channels composing each cluster. To ensure the quality of cluster signals, the cluster brain activity signal was computed for a participant if at least 3 channels with good quality signals were available.

### Analytic Plan

#### Preliminary Analyses: Quantification of synchrony

Cross-correlation within a maximum delay of 2 seconds (Azhari et al., 2020; Bizzego et al., 2019; Golland et al., 2014, 2015) between the normalized cluster signals of mother and child was computed, for each dyad, brain cluster and video.

This measure represents the synchrony of the brain activities of mother and child, while performing the task of watching the three videos together. We refer to this measure as *Copresence* synchrony.

To control for spurious correlations, we also computed the same measure from all possible mother-child pairs, except the true ones. We refer to this measure as *Control* synchrony; this measure will be influenced only by the effect of the audio-visual stimulation from the three videos.

*Copresence* and *Control* synchronies are compared with a Mann-Whitney test to ensure that *Copresence* synchrony is specific to the mother-child dyads, i.e. copresence has a significant effect in increasing the synchrony of mother-child brain activity. The computation of the synchrony measures was based on the *physynch* library (Bizzego, A., Gabrieli, G., Azhari, A., Setoh, P. and Esposito, G., 2019.).

#### Maternal Anxiety (MA) and Brain Synchrony

A Principal Component Analysis (PCA) with one component was adopted (based on *scikits-learn v0*.*22*.*1* Pedregosa,et al., (2011)) to derive a unique scale of Maternal Anxiety (MA) from the scores of the two questionnaires related to anxiety: Need for Approval (NFA) and Preoccupation (PREOP).

The estimated MA obtained from the PCA was negatively correlated with NFA (rs = - 0.86, p<0.001) and PREOP (rs=-0.79, p<0.001), i.e.: higher values were associated with reduced mother anxiety and vice versa. To facilitate the interpretation of the results, we inverted the estimated MA obtained from the PCA: the final MA estimate is positively associated with NFA and PREOP and, consequently, higher maternal anxiety (see Figure 4).

**Figure 4:**
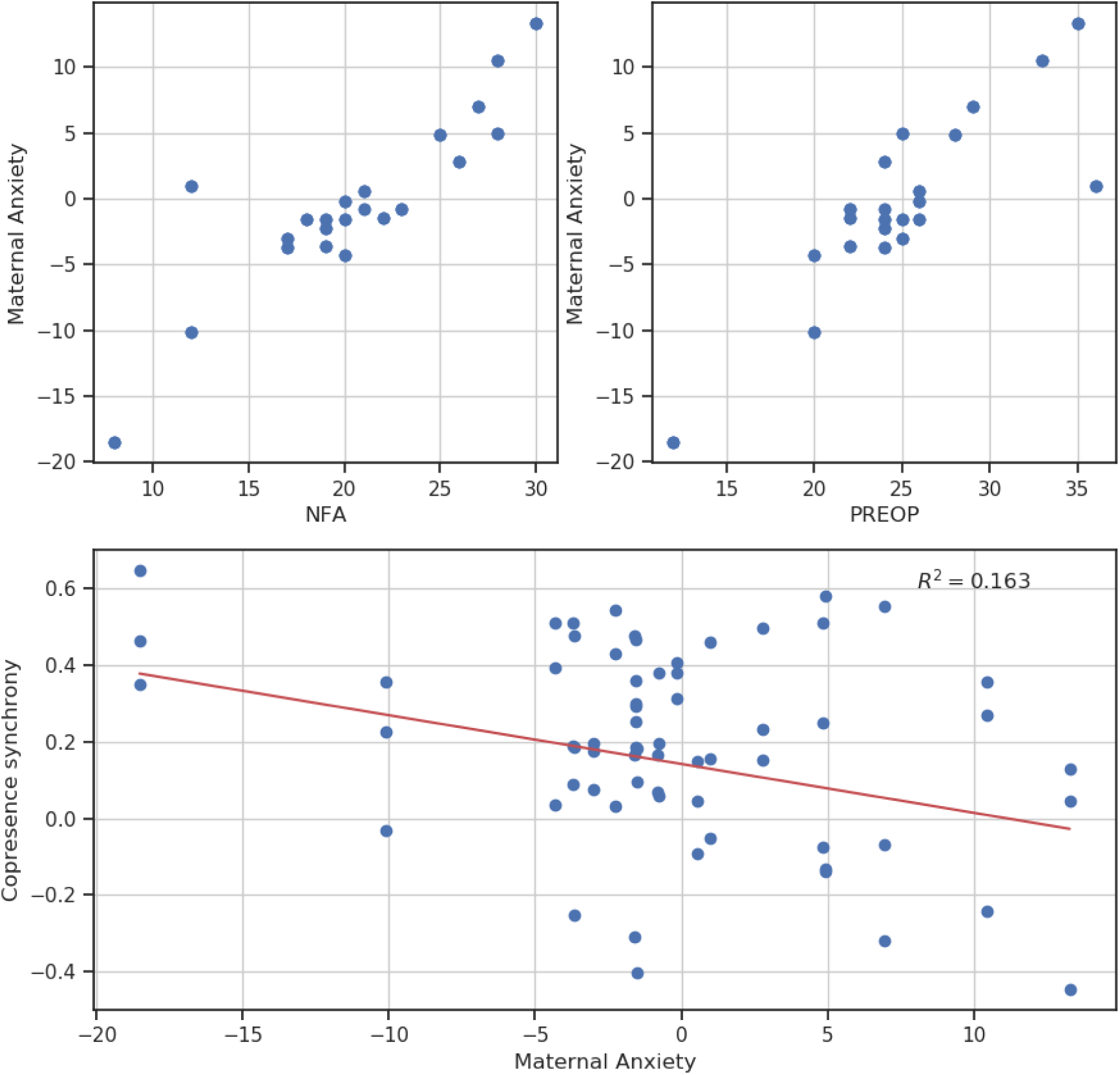
TOP: correlation between MA and the two anxiety scales. BOTTOM: correlation between MA and copresence synchrony.

The mean and standard deviation of synchrony values for each of the four clusters of copresence and control dyads are reported.

#### Inferential Analyses

The effect of Maternal Anxiety on the synchrony of mother-child brain activity was estimated by fitting a linear regression model on MA to predict copresence synchrony for each cluster. The linear regression model also included the audio-visual characteristics of the three videos (VC and AF0) as co-factors to account for the differences in stimulus properties across the videos that could potentially influence synchrony. In the regression analyses, False Discovery Rate (FDR) correction was applied to the four clusters.

## Results

### Preliminary Results

The Mann-Whitney test indicated that the measured synchrony was greater for true dyads (Median of copresence synchrony = 0.191) than surrogate dyads (Median of control synchrony = 0.162): U=763629, p=0.013, which means that copresence has a significant effect in modulating the synchrony of mother-child brain activity. The means and standard deviations of synchrony values for each of the 4 clusters of copresence and control dyads are reported in Table 2.

### Inferential Results

A significant linear association between copresence synchrony and MA was found for the medial right cluster (R^2^=0.163, p-corrected=0.029). As only the medial right cluster was found to be significantly correlated with MA, Spearman’s correlation test was only conducted for this cluster. In this cluster, the Spearman’s correlation between MA and copresence synchrony is rs = 0.251, p=0.043 (see Figure 4). No other cluster was significantly associated with MA (see Table 3).

**Table 3.** Results from inferential analyses of the correlation between Maternal Anxiety (MA) and mother-child brain synchrony.

| **Cluster** | **Coefficient** | **r<sup>2</sup>** | **p** | **p-corrected** |
| --- | --- | --- | --- | --- |
| FrontalLeft | 0.0044 | 0.019 | 0.33987 | 0.45315 |
| FrontalRight | -0.0024 | 0.026 | 0.63234 | 0.63234 |
| MedialLeft | -0.0096 | 0.076 | 0.09067 | 0.18134 |
| MedialRight | -0.0127 | 0.163 | 0.00717 | 0.02867 |

## Discussion

This study aimed to investigate whether maternal attachment style is associated with mother-child brain-to-brain synchrony when dyads engage in an everyday joint activity of watching animation shows together. We predicted that dyads with anxiously attached mothers would show less brain-to-brain synchrony. This hypothesis was fulfilled as dyads of mothers with higher Maternal Anxiety exhibited diminished brain-to-brain synchrony in the medial right cluster of the PFC. Reduced synchrony is observed in the medial right cluster of the prefrontal cortex, which overlaps with Brodmann Areas (BA) 8, 9, 45, and 46, where several stages of perceptual processing occur (Barbas & Zikopoulos, 2007; Medalla et al., 2007). The medial PFC, along with the temporo-parietal junction and the posterior cingulate cortex, have been established as the core seat of emotional (Atique et al., 2011; Corradi-Dell’Acqua et al., 2014) and cognitive mentalisation processes (Van Overwalle, 2009; Van Overwalle & Baetens, 2009). An anxiously-attached mother predominantly engages in emotional mentalisation, comprising automatic processes that rely on social cues (e.g., facial expressions) to form an understanding of the mental state of others (Fonagy & Luyten, 2009; Lieberman, 2007). She desires to be in close proximity with her child, leaving her greatly preoccupied with interpreting cues of whether the child is accepting of her (Mikulincer & Shaver, 2007; Feeney & Collins, 2001). This fixation with detecting the faintest signs of disapproval might detract her attention from the animation stimuli. While the child aligned his/her mental states to those of the characters in the videos screened to them, the anxiously-attached mother might not have fully engaged with the plots and characters of the animation shows, leading to a mismatch of brain signals in regions responsible for mentalisation processes between mother and child. As the medial PFC is an essential hub for mentalisation processes, brain-to-brain synchrony in this region reflects shared mutual attention and joint perceptual processing between individuals (Gvirts & Perlmutter, 2020). A lack of synchrony in the medial PFC between a mother with anxious-related attachment style and her child therefore suggests that the two of them differed in their intersubjective experiences during the activity, despite viewing the same animation stimuli together. It is likely that the mother’s constant need for confirmation of acceptance interfered with her ability to wholly engage in such bonding experiences. Counterintuitively, her preoccupation with the relationship could have undermined the quality of shared experiences that would have otherwise buttressed her feelings of social connectedness to her child. In the day-to-day context, the mother’s reduced attention during joint watching of television could hinder her ability to offer mental state comments in subsequent interactions with her child (e.g. reflecting the emotions of characters in the shows), which could have implications on the child’s development. Indeed, several studies have found that mother’s appropriate mental state comments predicted the child’s theory of mind ability in preschool (Laranjo et al., 2010; Meins et al., 2002). Future attachment research should investigate the potential for brain-to-brain synchrony to serve as a mechanism underlying parent-child bonding during joint passive activities, as well as uncover the association between dyadic brain synchrony and child development. This study had several limitations of which two are especially noteworthy. First, the scope of our investigation was limited to the prefrontal cortex. Several other regions of the brain which are connected to the lateral PFC might have exhibited similar patterns of synchrony in association with attachment anxiety. Future studies should extend the region of investigation to capture changes in synchrony in other areas of the brain that are implicated in mentalisation, including the subcortical structures like the basal ganglia and amygdala, and cortical areas such as the lateral temporal cortex (Fonagy & Luyten, 2009; Lieberman, 2007; Satpute & Lieberman, 2006). Second, our study only measured state anxiety in the context of attachment. Maternal trait anxiety could have also contributed to measured synchrony and should be investigated in future studies.

## Conclusion

Our study revealed that dyads with anxiously attached mothers exhibit less brain-to-brain synchrony in the medial right PFC when the pair engages in a typical joint activity of watching animation shows together. As this region is implicated in mentalisation, our findings suggest that, despite engaging in the same shared activity, an anxiously-attached mother’s preoccupation with social cues of affirmation or rejection from the child might give rise to differences in intersubjective experiences which interferes with the quality of bonding.

## Acknowledgements

This work was supported by XXX (G.E.) from XXX (G.E.), the XXX (M.H.B.), and an XXX (M.H.B.), funded by XXX (M.H.B.).

## References

Andrews-Hanna, J. R., Smallwood, J., & Spreng, R. N. (2014). The default network and self-generated thought: component processes, dynamic control, and clinical relevance. Annals of the New York Academy of Sciences, 1316, 29–52.

Atique, B., Erb, M., Gharabaghi, A., Grodd, W., & Anders, S. (2011). Task-specific activity and connectivity within the mentalizing network during emotion and intention mentalizing. NeuroImage, 55(4), 1899–1911.

Azhari, A., Leck, W. Q., Gabrieli, G., Bizzego, A., Rigo, P., Setoh, P., Bornstein, M. H., & Esposito, G. (2019a). Parenting Stress Undermines Mother-Child Brain-to-Brain Synchrony: A Hyperscanning Study. Scientific Reports, 9(1), 11407.

Azhari, A., Leck, W. Q., Gabrieli, G., Bizzego, A., Rigo, P., Setoh, P., Bornstein, M. H., & Esposito, G. (2019b). Parenting Stress Undermines Mother-Child Brain-to-Brain Synchrony: A Hyperscanning Study. Scientific Reports, 9(1), 11407.

Azhari, A., Lim, M., Bizzego, A., Gabrieli, G., Bornstein, M. H., & Esposito, G. (2020). Physical presence of spouse enhances brain-to-brain synchrony in co-parenting couples. Scientific Reports, 10(1), 7569.

Barbas, H., & Zikopoulos, B. (2007). The prefrontal cortex and flexible behavior. The Neuroscientist: A Review Journal Bringing Neurobiology, Neurology and Psychiatry, 13(5), 532–545.

Bartholomew, K. (1990). Avoidance of Intimacy: An Attachment Perspective. In Journal of Social and Personal Relationships (Vol. 7, Issue 2, pp. 147–178). https://doi.org/10.1177/0265407590072001

Bartholomew, K., & Horowitz, L. M. (1991). Attachment styles among young adults: A test of a four-category model. In Journal of Personality and Social Psychology (Vol. 61, Issue 2, pp. 226–244). https://doi.org/10.1037/0022-3514.61.2.226

Behrendt, H. F., Konrad, K., Perdue, K. L., & Firk, C. (2020). Infant brain responses to live face-to-face interaction with their mothers: Combining functional near-infrared spectroscopy (fNIRS) with a modified still-face paradigm. Infant Behavior & Development, 58, 101410.

Bizzego, A., Azhari, A., Campostrini, N., Truzzi, A., Ng, L. Y., Gabrieli, G., Bornstein, M. H., Setoh, P. &, Esposito, G. (2019). Strangers, Friends, and Lovers Show Different Physiological Synchrony in Different Emotional States. Behavioral Sciences, 10(1). https://doi.org/10.3390/bs10010011

Bizzego, A., Gabrieli, G., Azhari, A., Setoh, P. and Esposito, G. (n.d.). Computational methods for the assessment of empathic synchrony. In Esposito, A., Faundez-Zanuy, M., Morabito, F.C., Pasero, E. (Eds.) Smart Innovation Systems and Technologies: Progresses in Artificial Intelligence and Neural Systems, Springer.

Buehler, S. (2017). Handbook of Attachment, Third Edition: Theory, Research, and Clinical Applications. In Journal of Sex & Marital Therapy (Vol. 43, Issue 4, pp. 400–402). https://doi.org/10.1080/0092623x.2017.1317533

Corradi-Dell’Acqua, C., Hofstetter, C., & Vuilleumier, P. (2014). Cognitive and affective theory of mind share the same local patterns of activity in posterior temporal but not medial prefrontal cortex. In Social Cognitive and Affective Neuroscience (Vol. 9, Issue 8, pp. 1175–1184). https://doi.org/10.1093/scan/nst097

Crandell, L. E., Fitzgerald, H. E., & Whipple, E. E. (1997). Dyadic synchrony in parent–child interactions: A link with maternal representations of attachment relationships. In Infant Mental Health Journal (Vol. 18, Issue 3, pp. 247–264). https://doi.org/3.0.co;2-k”>10.1002/(sici)1097-0355(199723)18:3<247::aid-imhj2>3.0.co;2-k

Edelstein, R. S., Alexander, K. W., Shaver, P. R., Schaaf, J. M., Quas, J. A., Lovas, G. S., & Goodman, G. S. (2004). Adult attachment style and parental responsiveness during a stressful event. Attachment & Human Development, 6(1), 31–52.

Feeney, B. C., & Collins, N. L. (2001). Predictors of caregiving in adult intimate relationships: an attachment theoretical perspective. Journal of Personality and Social Psychology, 80(6), 972–994.

Feeney, J. A., & Noller, P. (1991). Attachment Style and Verbal Descriptions of Romantic Partners. In Journal of Social and Personal Relationships (Vol. 8, Issue 2, pp. 187–215). https://doi.org/10.1177/0265407591082003

Feeney, J. A., Noller, P., & Hanrahan, M. (2014). Attachment Style Questionnaire. In PsycTESTS Dataset. https://doi.org/10.1037/t29439-000

Feldman, R. (2007). On the origins of background emotions: from affect synchrony to symbolic expression. Emotion, 7(3), 601–611.

Feldman, R. (2012). Bio-behavioral Synchrony: A Model for Integrating Biological and Microsocial Behavioral Processes in the Study of Parenting. In Parenting (Vol. 12, Issues 2-3, pp. 154–164). https://doi.org/10.1080/15295192.2012.683342

Feldman, R. (2012). Interactive synchrony: A biobehavioral model of mutual influences in the formation of affiliative bonds in healthy and pathologicol development. In Neuropsychiatrie de l’Enfance et de l’Adolescence (Vol. 60, Issue 5, p. S2). https://doi.org/10.1016/j.neurenf.2012.04.016

Fleming, A. S., O’Day, D. H., & Kraemer, G. W. (1999). Neurobiology of mother–infant interactions: experience and central nervous system plasticity across development and generations. In Neuroscience & Biobehavioral Reviews (Vol. 23, Issue 5, pp. 673–685). https://doi.org/10.1016/s0149-7634(99)00011-1

Fonagy, P., & Luyten, P. (2009). A developmental, mentalization-based approach to the understanding and treatment of borderline personality disorder. Development and Psychopathology, 21(4), 1355–1381.

Golland, Y., Arzouan, Y., & Levit-Binnun, N. (2015). The Mere Co-Presence: Synchronization of Autonomic Signals and Emotional Responses across Co-Present Individuals Not Engaged in Direct Interaction. In PLOS ONE (Vol. 10, Issue 5, p. e0125804). https://doi.org/10.1371/journal.pone.0125804

Golland, Y., Keissar, K., & Levit-Binnun, N. (2014). Studying the dynamics of autonomic activity during emotional experience. Psychophysiology, 51(11), 1101–1111.

Gvirts, H. Z., & Perlmutter, R. (2020). What Guides Us to Neurally and Behaviorally Align With Anyone Specific? A Neurobiological Model Based on fNIRL Hyperscanning Studies. The Neuroscientist: A Review Journal Bringing Neurobiology, Neurology and Psychiatry, 26(2), 108–116.

Hasson, U., Nir, Y., Levy, I., Fuhrmann, G., & Malach, R. B. (2004). Intersubject synchronization of cortical activity during natural vision. Science, 303(5664), 1634–1640.

Hazan, C., & Shaver, P. (1987). Romantic love conceptualized as an attachment process. Journal Personality and Social Psychology, 52(3), 511–524.

Hopf, J.-M., & Mangun, G. R. (2000). Shifting visual attention in space: an electrophysiological analysis using high spatial resolution mapping. In Clinical Neurophysiology (Vol. 111, Issue 7, pp. 1241–1257). https://doi.org/10.1016/s1388-2457(00)00313-8

Jones, J. D., Brett, B. E., Ehrlich, K. B., Lejuez, C. W., & Cassidy, J. (2014). Maternal Attachment Style and Responses to Adolescents’ Negative Emotions: The Mediating Role of Maternal Emotion Regulation. In Parenting (Vol. 14, Issues 3-4, pp. 235–257). https://doi.org/10.1080/15295192.2014.972760

Janes, J. D., Chris Fraley, R., Ehrlich, K. B., Stern, J. A., Lejuez, C. W., Shaver, P. R., & Cassidy, J. (2018). Stability of Attachment Style in Adolescence: An Empirical Test of Alternative Developmental Processes. In Child Development (Vol. 89, Issue 3, pp. 871–880). https://doi.org/10.1111/cdev.12775

Laranjo, J., Bernier, A., Meins, E., & Carlson, S. M. (2010). Early Manifestations of Children’s Theory of Mind: The Roles of Maternal Mind-Mindedness and Infant Security of Attachment. In Infancy (Vol. 15, Issue 3, pp. 300–323). https://doi.org/10.1111/j.1532-7078.2009.00014.x

Licata, M., Zietlow, A.-L., Träuble, B., Sodian, B., & Reck, C. (2016a). Maternal Emotional Availability and Its Association with Maternal Psychopathology, Attachment Style Insecurity and Theory of Mind. Psychopathology, 49(5), 334–340.

Licata, M., Zietlow, A.-L., Träuble, B., Sodian, B., & Reck, C. (2016b). Maternal Emotional Availability and Its Association with Maternal Psychopathology, Attachment Style Insecurity and Theory of Mind. Psychopathology, 49(5), 334–340.

Lieberman, M. D. (2007). Social Cognitive Neuroscience: A Review of Core Processes. In Annual Review of Psychology (Vol. 58, Issue 1, pp. 259–289). https://doi.org/10.1146/annurev.psych.58.110405.085654

Syons, K. M., Stevenson, R. A., Owen, A. M., & Stojanoski, B. (n.d.). Examining the relationship between social cognition and neural synchrony during movies in children with and without autism. https://doi.org/10.1101/2020.03.06.981415

Maledalla, M., Lera, P., Feinberg, M., & Barbas, H. (2007). Specificity in inhibitory systems associated with prefrontal pathways to temporal cortex in primates. Cerebral Cortex, 17 Suppl 1, i136–i150.

Mofeins, E., Fernyhough, C., Wainwright, R., Gupta, M. D., Fradley, E., & Tuckey, M. (2002). Maternal Mind– Mindedness and Attachment Security as Predictors of Theory of Mind Understanding. In Child Development (Vol. 73, Issue 6, pp. 1715–1726). https://doi.org/10.1111/1467-8624.00501

Mikulincer, M., & Shaver, P. R. (2007). Attachment, group-related processes, and psychotherapy [Review of Attachment, group-related processes, and psychotherapy]. International Journal of Group Psychotherapy, 57(2), 233–245.

Mikulincer, M., & Shaver, P. R. (2010). Attachment Adulthood: Structure, Dynamics, and Change. Guilford Publications.

Nguyen, T., Schleihauf, H., Kayhan, E., Matthes, D., Vrtička, P., & Hoehl, S. (2020). The effects of interaction quality on neural synchrony during mother-child problem solving. Cortex; a Journal Devoted to the Study of the Nervous System and Behavior, 124, 235–249.

Pedregosa, F., Varoquaux, G., Gramfort, A., Michel, V., Thirion, B., Grisel, O., Blondel, M., Prettenhofer, P., Weiss, R., Dubourg, V, Vanderplas, J., Passos, A., Cournapeau, D., Brucher, M., Perrot, M. and Duchesnay, E. (2011). Scikit-learn: Machine Learning in Python. Journal of Machine Learning Research. JMLR, 12, 2825–2830.

Piazza, E. A., Hasenfratz, L., Hasson, U., & Lew-Williams, C. (2020). Infant and Adult Brains Are Coupled to the Dynamics of Natural Communication. Psychological Science, 31(1), 6–17.

Reindl, V., Gerloff, C., Scharke, W., & Konrad, K. (2018). Brain-to-brain synchrony in parent-child dyads and the relationship with emotion regulation revealed by fNIRS-based hyperscanning. In NeuroImage (Vol.178, pp. 493–502). https://doi.org/10.1016/j.neuroimage.2018.05.060

Rholes, W. S., Simpson, J. A., & Blakely, B. S. (1995). Adult attachment styles and mothers’ relationships with their young children. In Personal Relationships (Vol. 2, Issue 1, pp. 35–54). https://doi.org/10.1111/j.1475-6811.1995.tb00076.x

Satpute, A. B., & Lieberman, M. D. (2006). Integrating neurocognitive models of social cognition. Brain Research, 1079(1), 86–97.

Selcuk, E., Günaydin, G., Sumer, N., Harma, M., Salman, S., Hazan, C., Dogruyol, B., & Ozturk, A. (2010). Self-reported romantic attachment style predicts everyday maternal caregiving behavior at home. In Journal of Research in Personality (Vol. 44, Issue 4, pp. 544–549). https://doi.org/10.1016/j.jrp.2010.05.007

Sinhamay-Tsoory, S. G., & Aharon-Peretz, J. (2007). Dissociable prefrontal networks for cognitive and affective theory of mind: a lesion study. Neuropsychologia, 45(13), 3054–3067.

Sperling, M. B., & Berman, W. H. (1994). Attachment in Adults: Clinical and Developmental Perspectives. Guilford Press.

Stein, H., Dawn Koontz, A., Fonagy, P., Allen, J. G., Fultz, J., Brethour, J. R., Allen, D., & Evans, R. B. (2002). Adult attachment: What are the underlying dimensions? In Psychology and Psychotherapy: Theory, Research and Practice (Vol. 75, Issue 1, pp. 77–91). https://doi.org/10.1348/147608302169562

Tömasello, M. (1995). Joint attention as social cognition. In C. Moore & P. J. Dunham (Eds.) (Ed.), Joint attention: Its origins and role in development. (pp. 103–130). Lawrence Erlbaum Associates, Inc.

van IJzendoorn, M. H. (1995). Adult attachment representations, parental responsiveness, and infant attachment: a meta-analysis on the predictive validity of the Adult Attachment Interview. Psychological Bulletin, 117(3), 387–403.

Van Overwalle, F. (2009). Social cognition and the brain: A meta-analysis. In Human Brain Mapping (Vol. 30, Issue 3, pp. 829–858). https://doi.org/10.1002/hbm.20547

Van Overwalle, F., & Baetens, K. (2009). Understanding others’ actions and goals by mirror and mentalizing systems: a meta-analysis. NeuroImage, 48(3), 564–584.

Yeshurun, Y., Swanson, S., Simony, E., Chen, J., Lazaridi, automatic and controlled processes into C., Honey, C. J., & Hasson, U. (2017). Same Story, Different Story. In Psychological Science (Vol. 28, Issue 3, pp. 307–319). https://doi.org/10.1177/0956797616682029

