## Supplementary Material for "Maternal Anxious Attachment Style is Associated with Reduced Mother-Child Brain-to-Brain Synchrony During Passive TV Viewing"

### Supplementary Materials

#### *Analytic Plan: Beta-coefficient Analyses*

*Beta-coefficient Values*

At the level of within-subject analysis, haemodynamic response function (HRF) was specified and pre-whitening was omitted from the HbO signals. A convolution design matrix was used to plot the three stimuli against time axis to verify the correct order of stimuli presentation for each participant. Next, Discrete Cosine Transformation (DCT) temporal parameter with a high-pass period cut-off of 128 seconds and Gaussian Full Width at Half Maximum (FWHM) 4 models were applied. For each participant, a general linear model (GLM) was obtained from the HbO signals. From the GLM, the beta-coefficients of the stimuli were extracted and aggregated to form an average beta-coefficient for the participant. At the group-level, beta-coefficients of each participant were aggregated to form a GLM for group analyses. Beta-coefficients of channels that belong to the same cluster were averaged to generate a mean beta-coefficient for each cluster .

*Preliminary Analyses*

To investigate the effects of demographic and stimuli variables on brain responses, preliminary analyses were performed separately for mothers and children. For each cluster, three linear models were conducted where the child's gender, mother's age and video positivity ratings were inputted as factors against cluster beta-coefficient values. Video complexity, audio intensity and audio fundamentals were fitted as covariates across the models.

*Descriptive Analyses*

Beta-coefficient values were averaged across the four clusters for mothers and children. The mean and standard deviation of the mean beta-coefficients are reported.

*Inferential Analyses*

For each of the four clusters and each of the five subscales of attachment, a multiple linear regression model was performed with cluster beta-coefficient values as the dependent variables, attachment subscale scores as the independent variable and video complexity and audio fundamentals as controls. For example, when investigating the effect of Need for approval in the frontal right cluster, the model to be tested is: Beta (frontal right cluster) = Need for approval + (Video Complexity + Audio Fundamentals). Each set of regression analysis was conducted twice - separately for mothers and children. False Discovery Rate (FDR) correction was applied to correct p-values for multiple comparisons.

***Results: Beta-coefficient Analyses***

*Preliminary Results*

Similar to (Azhari et al., 2019), we performed three multiple linear regression models for mother's age, children's gender and video positivity rating. In each model, video complexity and audio fundamentals were fitted as co-variates. For mothers, none of the clusters, after p-value correction, was significantly correlated to the mother's age, children's gender and video positivity rating.

For children, a significant finding emerged in the frontal left cluster for mother's age ($R^2$= 0.05818, F(3,245) =5.045, p<0.01) and child's gender ($R^2$= 0.05816, F(3,245) =5.043, p<0.01), after p-value correction (Table 3a). However, closer examination of the frontal left cluster showed that the covariate video complexity was significant across all three regression models in this cluster ($R^2$=0.05816, F (3,246) =7.595, p<0.01). Since video complexity was controlled for throughout the study, only this variable was retained as our control for further analyses.

For children, a significant finding emerged in the frontal right cluster for video positivity ($R^2$= 0.02988, F (2,259)=3.988, p<0.05). Thus, video positivity was included in subsequent analyses with attachment subscales only for this cluster. Video complexity was also found to be significant in this cluster and was retained as a control in further analyses (Table 3b).

*Descriptive Results*

The mean and standard deviation of beta values for mother and child are reported in Table 4.

*Inferential Results*

Multiple linear regression analysis was conducted to examine if any attachment subscale significantly predicted cluster activation in the PFC for both the mother and the child. For both mothers and children, no significant relationship was found between any attachment subscale and PFC cluster activations after correction.

***Results: Synchrony Analyses (Insignificant Findings)***

*Confidence* was not significantly associated with distance index in the frontal left ($R^2$= -0.00115, F(3, 220)=0.915, p>0.05), frontal right ($R^2$=-0.00686, F(3, 225)=0.482, p>0.05), medial left ($R^2$= 0.0141, F(3, 299)=2.438, p>0.05) and medial right ($R^2$=-0.00391, F(3, 255)=0.665, p>0.05) clusters.

*Discomfort with closeness* was not significantly associated with distance index in the frontal left ($R^2$= 0.00208, F(3, 220)=1.155, p>0.05), frontal right ($R^2$=-0.0045, F(3, 225)=0.66, p>0.05), medial left ($R^2$= 0.00346, F(3, 299)=1.349, p>0.05) and medial right ($R^2$=0.00335, F(3, 255)=1.289, p>0.05) clusters.

*Relationships as Secondary* was not significantly associated with distance index in the frontal left ($R^2$= 0.000866, F(3, 220)=1.064, p>0.05), frontal right ($R^2$=-0.002, F(3, 225)=0.848, p>0.05), medial left ($R^2$= 0.00519, F(3, 299)=1.525, p>0.05) and medial right ($R^2$= -0.00381, F(3, 255)=0.673, p>0.05) clusters.

*Preoccupation with relationships* was not significantly associated with distance index in the frontal left ($R^2$= 0.00148, F(3, 220)=1.11, p>0.05), frontal right ($R^2$= -0.00824, F(3, 225)=0.379, p>0.05), medial left ($R^2$= 0.00394, F(3, 299)=1.398, p>0.05) and medial right ($R^2$=-0.00482, F(3, 255)=0.588, p>0.05) clusters.

Table 3a. Frontal left cluster of children: Beta coefficients of video complexity for regression models with mother's age, children's gender and positivity ratings.

| Models | *b* | *SE* | *t* | *p* |
|---|---|---|---|---|
| Mother's Age | -1.04e-09 | 2.68e-10 | -3.87 | 0.000139 |
| Children's Gender | -1.04e-09 | 2.68e-10 | -3.87 | 0.000139 |
| Video Positivity Rating | -1.14e-09 | 3.02e-10 | -3.79 | 0.000190 |

Table 3b. Frontal right cluster of children: Beta coefficients of video positivity ratings and video complexity covariates. Values based on preliminary analyses in Azhari et al. (2019).

| Variable | *b* | *SE* | *t* | *p* |
|---|---|---|---|---|
| Video Positivity Rating | - 3.24e-06 | 1.40e-06 | 2.31 | 0.0214 |
| Video Complexity | -8.88e-10 | 3.49e-10 | -2.55 | 0.0114 |

Table 4. Mean and standard deviation of the beta values of the clusters for mothers and children.

| Clusters | Mean beta values (Mothers) | Standard Deviation (Mothers) | Mean beta values (Children) | Standard Deviation (Children) |
|---|---|---|---|---|
| Frontal left | -5.25E-07 | 2.11E-06 | -2.71E-06 | 2.74E-06 |
| Frontal right | 8.82E-07 | 2.92E-06 | -1.02E-06 | 1.22E-06 |
| Medial left | 1.07E-06 | 6.35E-06 | -2.79E-06 | 2.62E-06 |
| Medial right | 4.09E-06 | 6.971E-06 | -1.55E-06 | 5.00E-06 |
